# Modeling and optimization of a central diamond shape threefold hexagon metamaterial sensor for glioblastoma cell detection

**DOI:** 10.64898/2026.03.11.710974

**Authors:** M.R. Foysal, Brinta Dey, M. Ahmed, Laboni Keya, S.M.A. Haque

## Abstract

The present study introduces a novel design and analysis of a sensor based on a terahertz metamaterial absorber (MMA) to identify Glioblastoma cell by employing microwave imaging techniques. Terahertz (THz) frequencies offer unique advantages for biomedical applications. Computer Simulation Technology (CST) employs a finite integration (FIT) approach to simulate the suggested structure in the resonant frequency (RF) range of 4.5 THz to 6 THz. The crystal structure displays three distinct absorption peaks at resonance frequencies. The MTA can absorb energy in three specific spectral bands: 4.782 THz, 5.30 THz, and 5.7319 THz. At these frequencies, the MTM achieves exceptionally high absorption, reaching 99.99%, 99.98%, and 99.68% peak absorption, respectively. The electric field (E), magnetic field (H), and surface current of the MTM are also examined. Finally, detecting Glioblastoma cells is also being investigated by analyzing the E-field H-field using microwave imaging. The suggested biosensor features a high-quality factor of 143.63, a frequency shifts per refractive index of 1.45 THz/RIU. In this study, the PCR value is measured at 0.95 at a frequency of 4.782 THz, indicating a high efficiency in polarization conversion. Polarization Conversion Ratio (PCR) quantifies the efficiency of a metamaterial in converting the polarization of an incident electromagnetic wave. Extensive simulation studies confirm the sensor’s capability to distinguish between healthy and cancerous cervical tissue. The suggested MMA-based sensor has numerous advantages and can be utilized for Glioblastoma cell detection.

## INTRODUCTION

Metamaterials (MTMs) are artificially engineered materials with unique electromagnetic properties, such as negative permittivity and permeability, complete absorption [1], complete transmission, and perfect lensing, invisibility [2], negative refractive index [3] which are not found in nature. Due to their potential applications in optics, biomedical imaging and sensing, liquid sensing [4], electronics, photovoltaic devices, solar cells, super lenses and cloaking these exceptional characteristics have scintillated significant research interest. One of the most prolific applications of MTMs, particularly in the terahertz (THz) regime is sensing technologies, where they enable highly sensitive detection of micro-scale substances. THz waves, spanning frequencies between 0.1 and 10 THz, occupy an intermediate region in the electromagnetic spectrum between infrared and microwave radiation [5]. Unlike X-rays, THz waves are non-ionizing and can interact with weak molecular forces, such as hydrogen bonds and van der Waals interactions, making them highly effective for identifying macromolecules [6]. Given their diverse capabilities, MTMs and their derivatives, such as Metamaterial Absorbers (MMAs), have become exigent in modern research, particularly in biomedical sensing and imaging applications.

MMAs are specially designed MTM structures that can generate intensified electromagnetic fields in localized regions. This improves the interaction between THz vibrations and the surrounding material, hence raising detection sensitivity. Because of a small variation in the refractive index (RI) of a substance applied to the MMA surface can change the surrounding permittivity, leading to a shift in the resonance dip of the MMA’s optical response. MMA-based sensors can effectively detect and characterize substances by analyzing these shifts in the absorption spectrum. This sensing technique, which leverages RI sensitivity, has exhibited immense potential in biomedical applications, including different kinds of cell detection. The ability of MMAs to enhance THz wave interactions with biomolecules offers a promising approach for non-invasive and highly sensitive medical diagnostics. Thus, the development and enhancement of MTM based RI sensors remain crucial in driving next generation sensing technologies forward.

Since many relevant works exist about the application of metamaterial absorbers, this section describes some of them, primarily focusing on biosensing and other applications. N. Hamza et al. [7] initiated an innovative concept for Non-melanoma skin cancer detection. The sensor exhibits an overall absorptivity of 99.5% at 0.904 THz, accompanied by a QF of 12.8, 13.5 and a resonance shift of 0.0515, 0.076 THz per refractive index unit (RIU). Tan et al. [8] developed a biosensor for breast cancer diagnostics using a terahertz-graphene metasurface. This simulation illustrates that the performance parameters such as the S, FOM, and QF for detecting a breast cell are 1.21 THz/RIU, 2.75 (1/RIU), and 2.43, respectively. Yang et al. [9] suggested a single and dual-resonance characteristic specifically suited for Gas, environmental and biomedical sensors. The sensor’s parameters, such as the FOM, S and QF, are calculated to be 50.7, 40 (1/RIU), 0.54, 1.12 (THz/RIU) and 57.4, 44.7. Jain et al. [10] introduced machine learning assisted hepta band THz metamaterial absorber for biomedical applications. This programmable detector has quality attributes of 117, resonance shift per RI of 4.72 THz, and FOM of 44 (1/RIU) respectively. Abdulkarim et al. [11] proposed an Highly Sensitive Dual-Band Terahertz Metamaterial Absorber for Biomedical Applications to achieve biomedical application. The computational study indicates that the relative sensitivity of the two resonance peaks of the structure is 0.0968 THz/RIU and 0.1182 THz/RIU. The simmulation exhibits a QF and FOM of 126 and 50.7 (1/RIU) respectively.

The present study presents a novel MMA structure capable of detecting glioblastoma cell. The method which is used in fabrication involves a gold resonator, a PTFE substrate, and a gold ground layer. The novelty of this research lies within the sensor’s excellent PCR value, sensitivity, and high Q factor, exceeding the cited literature. The detector operates by measuring the variation in refractive index in the analyte layer and comparing healthy and glioblastoma cell based on the shift in the absorption curve.

## METHODOLOGY AND UNIT CELL DESIGN

The proposed metamaterial absorber (MMA) consists of three distinct layers: a resonator as the uppermost layer, a dielectric medium in the middle, and a metallic substrate at the bottom. The resonator film and the ground plane are positioned on opposite sides, separated by the dielectric spacer. Gold (Au) is utilized for both the resonator and substrate due to its superior electrical conductivity (σ = 4.561 × 10^7^ S/m) and thermal conductivity (k = 314.0 W/m·K). These properties facilitate resonance within the targeted frequency range by enabling the generation of inductance and capacitance [12] . Additionally, gold exhibits notable physical characteristics, including a specific heat capacity (C□) of 0.13 J/kg·K, a density of 19,320 kg/m^3^, a Young’s modulus of 78 GPa, and a Poisson’s ratio of 0.42.

Teflon (PTFE) is selected as the dielectric material due to its low relative permittivity and minimal loss tangent, making it well-suited for terahertz (THz) absorption applications. Furthermore, PTFE demonstrates remarkable thermal resistance and significant flexural strength. The MMA is designed to interact with a THz Transverse Electric (TE) wave, where both the polarization angle and incidence angle are set to zero degrees. The dielectric properties of PTFE include a permittivity of 2.1 and a permeability of 1, while its Poisson’s ratio is 0.4. Additionally, PTFE has a thermal conductivity of k = 0.2 W/m·K.

The structural design of the MMA is symmetric, as depicted in Fig. 1(a), ensuring polarization insensitivity and achieving an almost negligible polarization conversion ratio (PCR). The symmetry enhances the absorber’s performance by maintaining consistent response characteristics. The unit cell dimensions are 120 × 120 μm^2^, with individual layer thicknesses of 2 μm for the ground plane, 15 μm for the dielectric substrate, and 1 μm for the resonator. A summary of the key parameters employed in the design is presented in Table 1.

**Fig 1.**
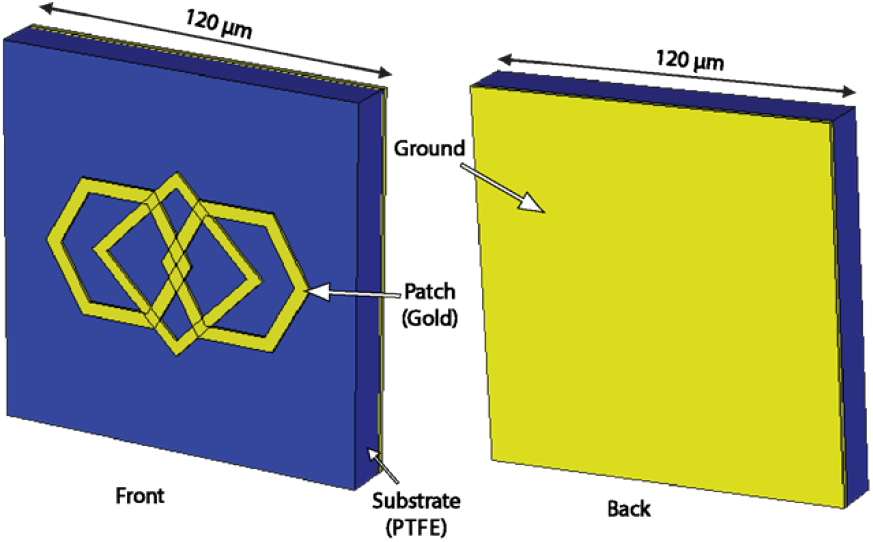
Perspective view of the designed material.

**Table 1:**
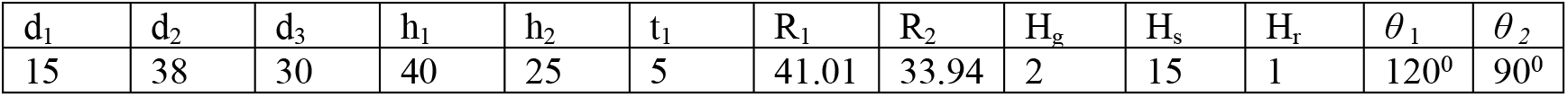
Necessary Parametric Values of the suggested model:

| $d_1$ | $d_2$ | $d_3$ | $h_1$ | $h_2$ | $t_1$ | $R_1$ | $R_2$ | $H_g$ | $H_s$ | $H_r$ | $\theta_1$ | $\theta_2$ |
| --- | --- | --- | --- | --- | --- | --- | --- | --- | --- | --- | --- | --- |
| 15 | 38 | 30 | 40 | 25 | 5 | 41.01 | 33.94 | 2 | 15 | 1 | $120^\circ$ | $90^\circ$ |

**Table 2.** Comparison table of MTM-based biosensors of different applications.

| Ref. | Techniques used | Operating Frequency (THz) | Q | S (THz/RIU) | Applications | Published in | Year of publication |
| --- | --- | --- | --- | --- | --- | --- | --- |
| [8] | Al/PET/Al | 0–1.2 | 12.8, 13.5 | 0.0515, 0.076 | Non-melanoma skin cancer detection. | IEEE Access | 2023 |
| [10] | Au/Poli-Si/Si<sub>3</sub>N<sub>4</sub>/Quartz | 0.2–20 | 57.4, 44.7 | 0.54, 1.12 | Gas, environmental and biomedical sensors. | Nanomaterials | 2021 |
| [22] | SiO<sub>2</sub>/Ion-gel/Air/Au/Graphene | 0.1–20 | – | 0.45, 0.717 | Analyte sensing | Frontiers in Physics | 2022 |
| [9] | Graphene/SiO<sub>2</sub> | 0.5–2.5 | 2.43 | 1.21 | Breast cancer detection | Nanomaterials | 2022 |
| [11] | Au/Polyimide/Au | 1–8.5 | 117 | 4.72 | Analyte sensing | Scientific Reports | 2023 |
| [23] | Au/GaAs/Au | 0–4 | – | 0.103, 0.095 | Microorganism detection | Scientific Reports | 2023 |
| [12] | Al/Photoresist/ | 0–1 | 70, 126 | 0.0968, 0.1182 | Biomedical application | American | 2022 |
|  | PET/Quartz |  |  |  |  | Chemical Society |  |
| [24] | Graphene/SiO<sub>2</sub>/Au | 2–6 | 13.76 | 0.851 | RI biosensor | Research Square | 2021 |
| **Work** | **Au/PTFE (Teflon)/Au** | **4–6** | **143.63** | **1.45** | **Early-stage brain cancer detection** |  | **2025** |

In terahertz (THz) applications, the transmission of electromagnetic waves is typically obstructed when the thickness of the metallic ground layer exceeds the skin depth or penetration depth of the THz wave. Given that the bottom metal plate completely blocks THz transmission, further increasing its thickness is unlikely to significantly impact the absorption spectrum. This is because transmission has already been reduced to zero, meaning that additional thickness would not influence the absorber’s overall performance. Consequently, in principle, as long as the metal layer is sufficiently thick to prevent transmission, variations in its thickness should not alter the absorption characteristics of the structure.

To ensure the validity and accuracy of the simulation results for the proposed structure, it is essential to analyze its behavior under different boundary conditions. The reliability and consistency of the numerical findings can be confirmed by comparing them with analytical solutions, experimental measurements, or simulations performed using alternative computational tools. All numerical analyses of the proposed design were conducted using CST Studio Suite, which employs the Finite Integration Technique (FIT) [13].

For the Transverse Electromagnetic (TEM) mode, the boundary conditions are defined such that a perfect electric conductor (PEC) is applied along the y-z plane, a perfect magnetic conductor (PMC) is assigned to the x-z plane, and the x-y plane remains open to allow THz wave propagation. In the case of the Transverse Electric (TE) and Transverse Magnetic (TM) modes, the simulations incorporate a Floquet port in the z-direction and implement a master-slave boundary condition along the x and y directions. The unit cell configuration constrains the x and y directions, while an open boundary is applied along the z-axis to facilitate wave propagation. The boundary conditions for the metamaterial absorber are illustrated in Fig. 2.

**Fig 2.**
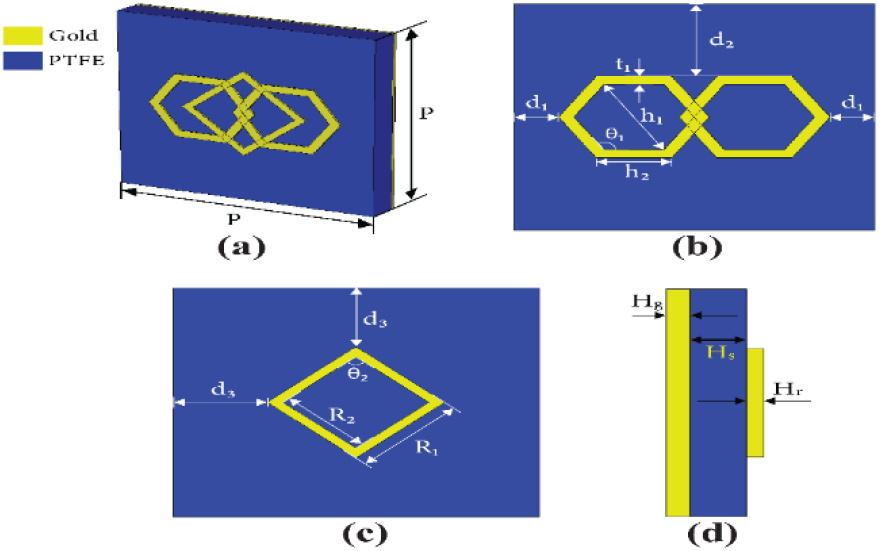
The detailed dimension of the unit cell.

**Fig 3.**
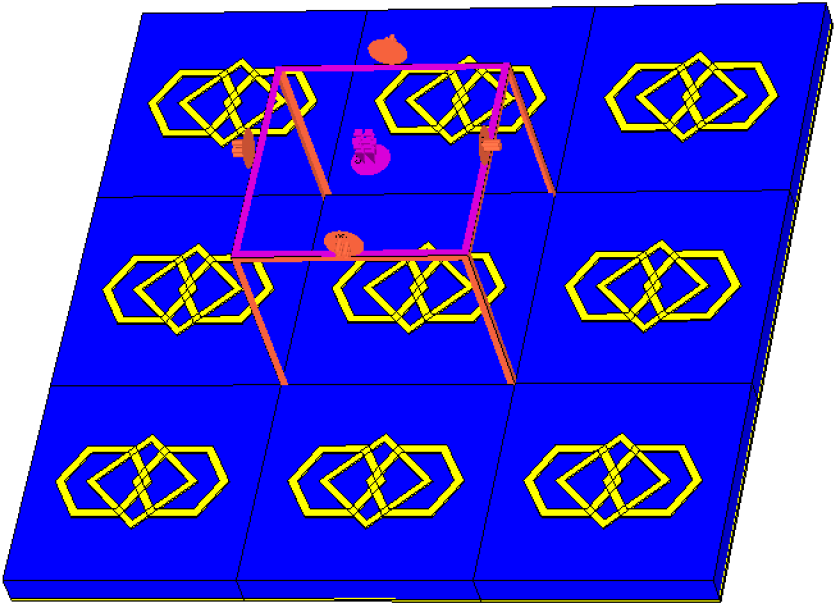
MMA with boundary conditions.

**Fig 4.**
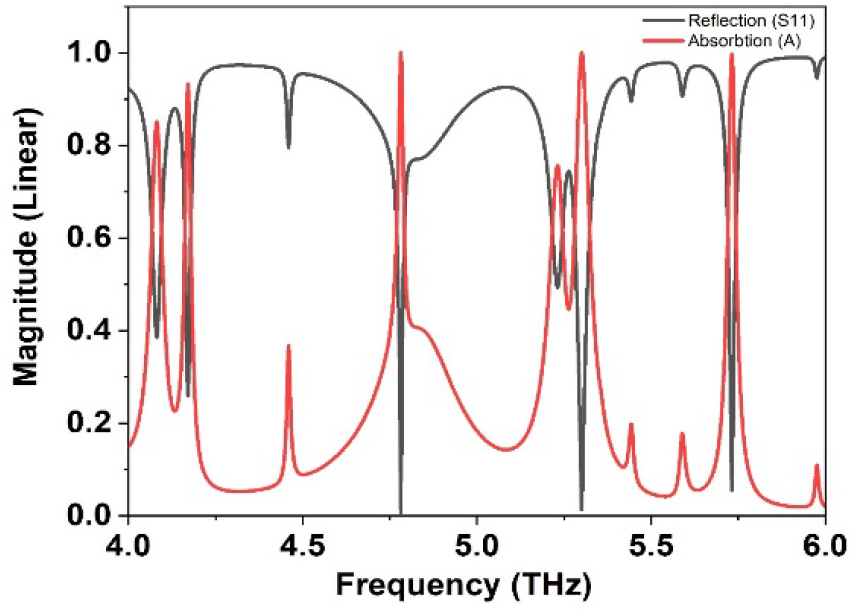
S_11_ and absorption of the suggested model.

**Fig 5.**
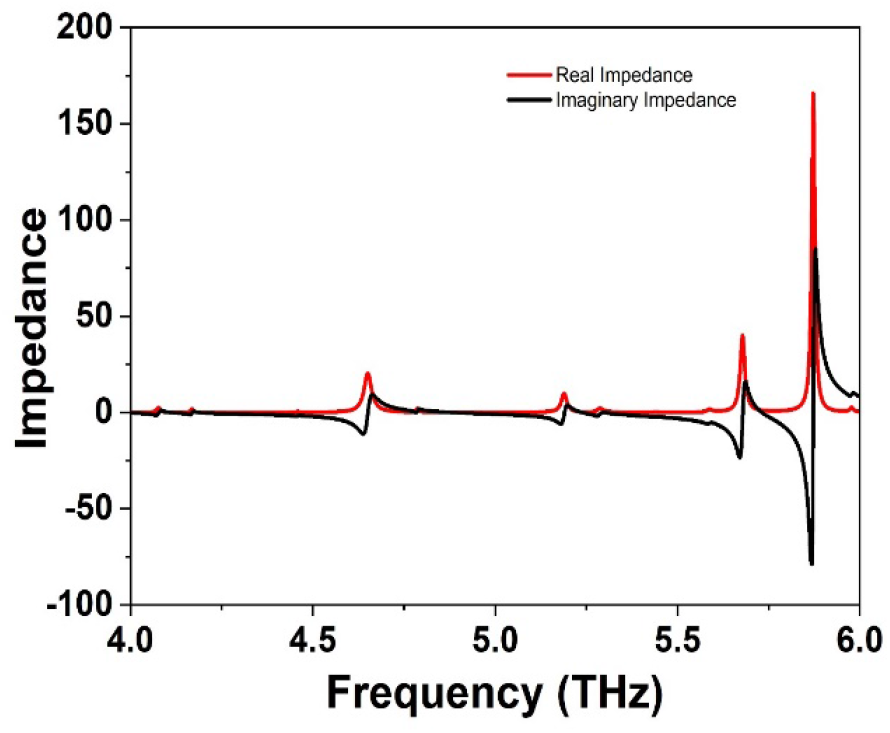
Impedance of the proposed MMA.

The wave interaction with the structure can be interpreted using Snell’s law. When a wave transitions from a right-handed material to a medium with a negative refractive index, the reflected wave remains within the same region as the incident wave, as depicted in Fig. 2. This phenomenon is governed by the conservation of the tangential components of both incident and refracted wave vectors, as well as the causality principle, which dictates that energy must propagate away from the interface. The proposed structure is designed to operate under a THz TE wave with a polarization angle (ϕ) of 0° and normal incidence (θ) of 0°. The wave propagates along the z-direction, featuring a vertically polarized electric field (E_x_) and a horizontally polarized magnetic field (H_γ_).

## RESULTS AND DISCUSSIONS

The effectiveness of the metamaterial sensor is heavily influenced by the absorption within the specified frequency range. CST is utilized to derive performance characteristics like s-parameters and absorption coefficients through the FIT. The MMA has a ground of solid copper; thus, there will be very little transmission through it. Therefore, absorption may be calculated by determining the coefficients of reflection and transmission using the following equation at 1 [14]:

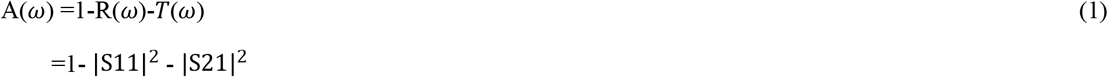

Where, R(*ω*) = |S11|^2^ is the reflection coefficient and the transmission coefficient is denoted as T(*ω*) = |S21|^2^ as the copper ground restricts the transmission of the wave, the transmission coefficient becomes zero. Thus equation 1 becomes:

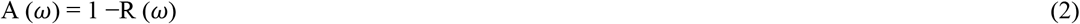

At 4.408, 6.824, and 11.048 GHz, this design shows three resonance peaks of absorption: 91%, 99.74%, and 99.99%, respectively. Since the bottom copper essentially prevented incoming radiation from being transmitted, S21 was predicted to be zero. The observation revealed that the proposed MMA had DNG features, or negative mu and negative epsilon attributes.

The reflection coefficient of the designed MMA can be written as[15].

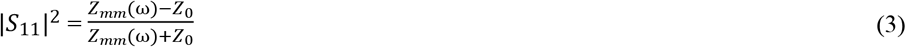

*Z*_mm_ represents the resistance of the metamaterial absorber (MMA), while *Z*_0_ signifies the resistance in an open space environment. Understanding *Z*_*mm*_ helps us comprehend the unique properties of the absorber and how it interacts with electromagnetic waves. This is essential for improving the effectiveness of various applications [16].

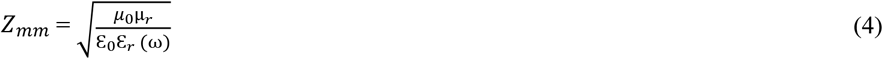

The factors, permeability and permittivity of vacuum space are represented by μ_0_ and ε_0_, respectively, while the frequency-lean on relative permeability and relative permittivity are denoted by μ_r_(ω) and ε_r_(ω), respectively. If Z_0_, the intrinsic impedance is equal to *Z*_*mm*_ in (empty space), then the incident electromagnetic wave will not be reflected from the absorber [17].

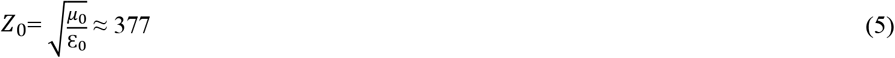

Where *µ*_0_ is the permeability of vacuum space and ε_0_ is the permittivity of vacuum space, gives the intrinsic impedance of free space. The value of ≈377Ω represents the rate of the E-field to the H-field in an electromagnetic wave traveling through a vacuum. The close approximation of 377 ohms reflects the physical relationship between the fundamental constants of nature, linking the behavior of electric and magnetic fields in free space.

### Optimization of Reflection (S11) and Absorption (A)

The graph illustrates the reflection (S11) and absorption (A) characteristics of a material across a frequency range of 4.0 to 6.0 THz. The reflection magnitude, represented by S11, shows a decreasing trend as the frequency increases, indicating that the material reflects less energy at higher frequencies. Conversely, the absorption (A) exhibits an increasing trend, suggesting that the material absorbs more energy as the frequency rises. Notably, the reflection spectrum displays several sharp dips, indicating strong absorption at specific frequencies. For instance, at approximately 4.2 THz, the reflection drops to around 0.1, suggesting a corresponding absorption peak near 0.9. Similarly, at 4.7 THz, the reflection reaches a minimum of approximately 0.05, indicating a high absorption of around 0.95. Conversely, at 5.0 THz, the reflection peaks near 0.9, implying minimal absorption. These fluctuations suggest the material’s selective interaction with incident radiation, potentially due to resonant phenomena within its structure. This behavior is typical in materials where higher frequencies lead to greater interaction with the material’s internal structure, resulting in enhanced absorption and reduced reflection. The data is crucial for applications in designing materials for electromagnetic shielding or optimizing performance in terahertz technologies.

### Optimization of Impedance

The graph illustrates the real and imaginary components of impedance as a function of frequency, spanning the terahertz (THz) range from 4.0 to 6.0 THz. The real impedance, representing the resistive part, shows fluctuations with several sharp peaks, indicating resonances at specific frequencies.

The imaginary part of the equivalent impedance is almost 0 at resonance frequencies, whereas the real part is almost one. This means that there are less reflections at the resonance frequency due to superior impedance adaptation within the absorber and the surrounding free space. The net impedance of the absorber can be computed using the S-parameters and the following formula:

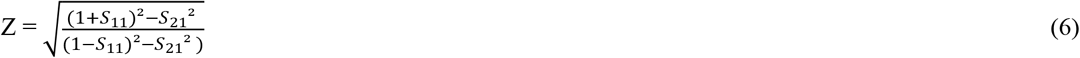

For instance, at approximately 5.8 THz, the real impedance surges to over 150 ohms. Concurrently, the imaginary impedance, representing the reactive part, exhibits corresponding sharp transitions, crossing zero at these resonant frequencies and indicating a shift from capacitive to inductive behavior, or vice versa. These fluctuations in both real and imaginary impedance signify the material’s dynamic response to varying frequencies, potentially revealing information about its intrinsic electrical properties and resonant modes. This behavior is critical for understanding the material’s response to terahertz radiation, which is essential for applications in telecommunications, sensing, and material characterization. The data highlights the frequency-dependent nature of impedance, providing insights into the material’s electromagnetic properties and potential uses in advanced technologies. The observed peaks and dips in impedance may correspond to resonant frequencies where the material’s interaction with electromagnetic waves is maximized or minimized, further emphasizing the importance of these measurements in optimizing material performance for specific applications.

### Optimization of PCR

The PCR quantifies the effectiveness of a metamaterial or metasurface in transforming the polarization of an incoming electromagnetic wave. The PCR value of the suggested MMA is calculated to confirm that it functions as an absorber rather than a polarization converter. The PCR value must be close to zero to verify the insensitivity of the suggested meta-structure unit cell to polarization conversion.

The graph represents the Polarization Conversion Ratio (PCR) as a function of frequency, ranging from 4.0 to 6.0 THz. The PCR values fluctuate between 0.0 and 1.0, indicating variations in the efficiency of polarization conversion at different frequencies. The peaks and troughs in the graph suggest that the material or device exhibits optimal polarization conversion at specific resonant frequencies, while efficiency drops at others. This frequency-dependent behavior is crucial for applications in terahertz technology, such as energy harvesting, signal processing, and communication systems.

### Optimization of Permittivity and permeability

The image depicts a graph illustrating the permittivity of a material as a function of frequency, with the x-axis representing frequency in terahertz (THz) ranging from 4.0 to 6.0 THz, and the y-axis representing permittivity values divided into real and imaginary parts. The real part of permittivity, which indicates the material’s ability to store electrical energy in an electric field, ranges approximately from -400 to 800, while the imaginary part, representing energy loss or dissipation within the material, spans a similar range. This graph is essential for analyzing the dielectric properties of materials, as it shows how both components of permittivity vary with frequency, providing valuable insights into the material’s response to electromagnetic fields.

The image displays a graph illustrating the permeability of a material as a function of frequency, with the x-axis representing frequency in terahertz (THz) ranging from 4.0 to 6.0 THz, and the y-axis representing permeability values divided into real and imaginary parts. The real part of permeability, which indicates the material’s ability to support the formation of a magnetic field, ranges approximately from -20000 to 40000, while the imaginary part, representing magnetic losses or energy dissipation within the material, spans a similar range. In this particular case, the permittivity shows a negative trend, while the permeability also remains negative. Based on the Fig. 6 and Fig. 7 the observed Metamaterial Absorber (MMA) demonstrated the characteristics of a double-negative metamaterial, exhibiting both negative permittivity and permeability [18].

**Fig. 6.**
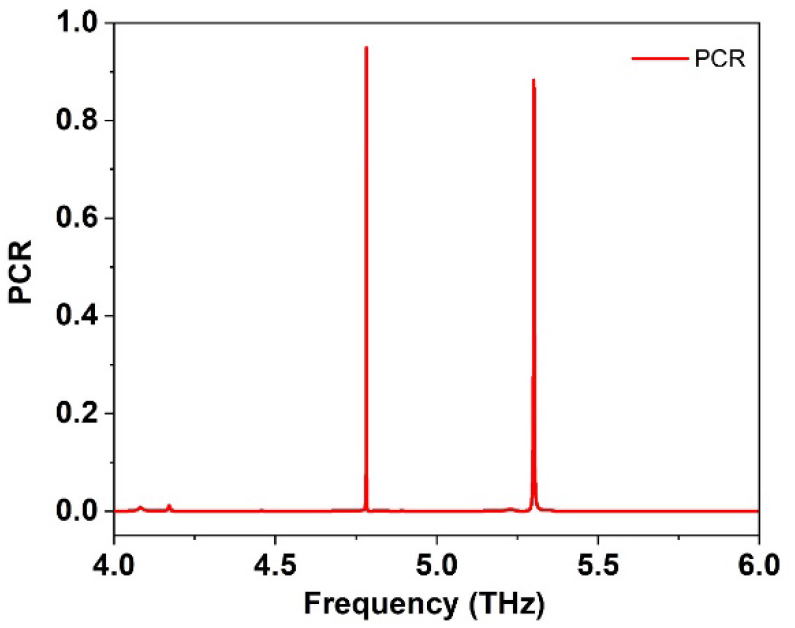
PCR of the proposed MMA.

**Fig 7.**
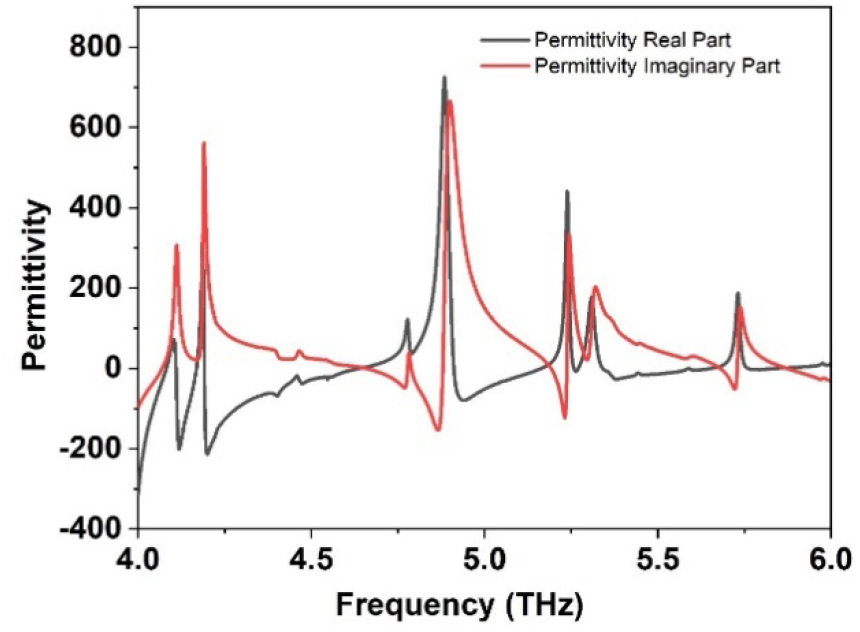
Permittivity of the proposed MMA.

**Fig 8.**
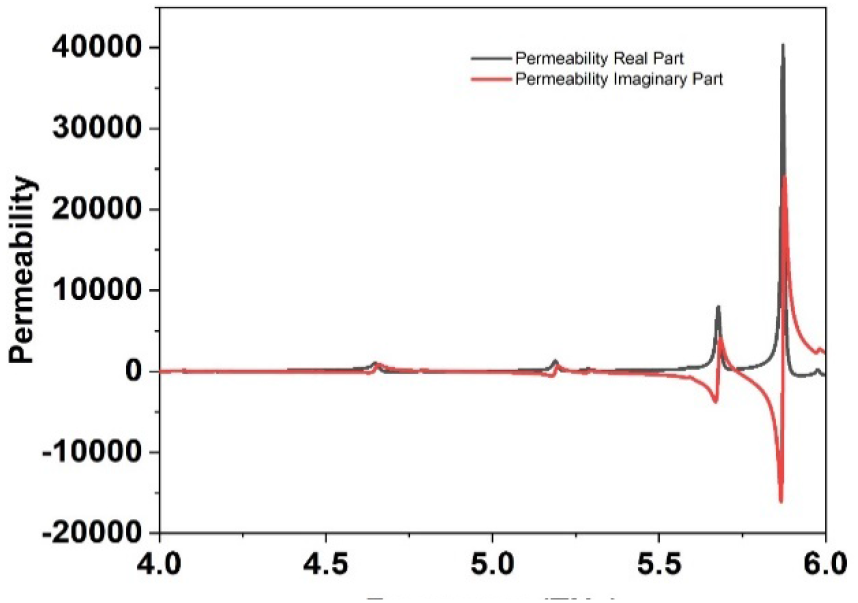
Permeability of the suggested absorber.

**Fig 9.**
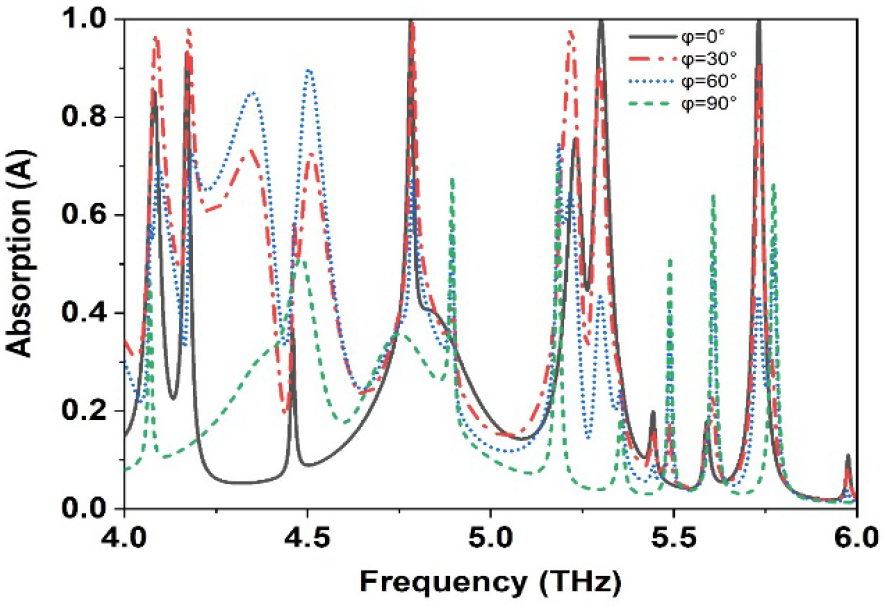
Variation of absorption the of polarization angle of the proposed MMA.

**Fig 10.**
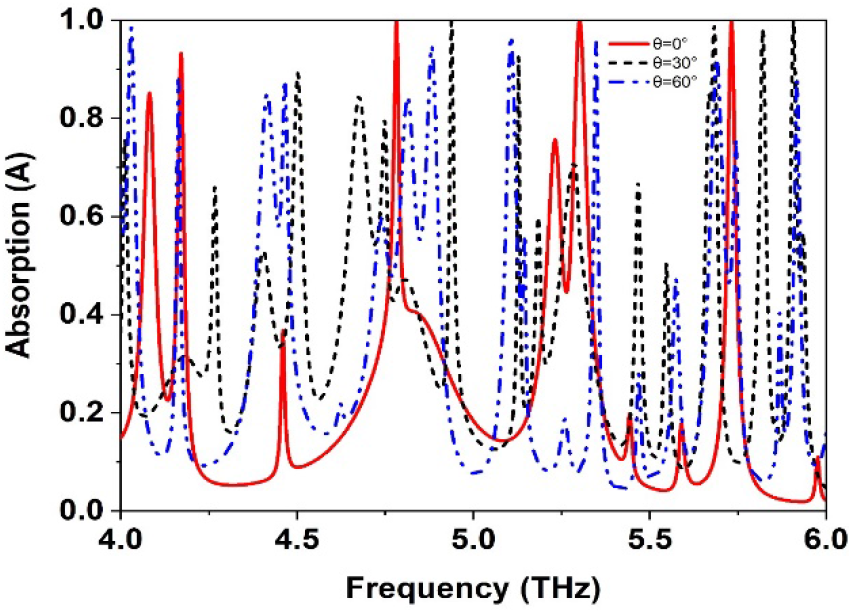
Variation of absorption the of incident angle of the proposed MMA.

### Optimization of the variation of polarization angle and incident angle

The image depicts a graph illustrating the absorption characteristics of a material as a function of frequency, with a focus on the polarization angle. The x-axis represents frequency in terahertz (THz), ranging from 4.0 to 6.0 THz, while the y-axis represents absorption (A), ranging from 0.0 (no absorption) to 1.0 (complete absorption). The graph shows how absorption varies with frequency for different polarization angles, specifically at 0°, 30°, 60°, and 90°, across the frequency range. Sharp peaks, known as resonances, indicate strong absorption at specific frequencies, highlighting the material’s significant interaction with incoming radiation.

This data is essential for understanding the interaction of electromagnetic waves with materials, particularly in fields like optics, materials science, and photonics. The variations in absorption with frequency and polarization provide insights into the material’s response to different light polarizations, which is critical for applications in optical devices, sensors, and communication technologies. Analyzing absorption at various polarization angles helps optimize the material’s performance for applications where polarization sensitivity is key.

This graph illustrates the complex interplay between frequency, incident angle, and absorption in a material or structure within the terahertz (THz) regime. The absorption spectra exhibit a pronounced oscillatory behavior, revealing multiple resonant modes that are sensitive to the angle of incidence (θ). As θ increases from 0° to 60°, significant shifts in peak positions and intensities occur, indicating changes in the effective optical path length and mode coupling. This angle-dependent absorption suggests the potential for tuning the optical response, which could be exploited in applications such as sensing, filtering, and modulating THz radiation.

### Optimization of Electric-field, Magnetic-field, and Surface current distribution

Figure11 serves as a crucial resource for understanding the behavior of MMA by illustrating the distribution of surface currents, electric fields, and magnetic fields. The analysis of this figure indicates that the magnetic field exhibits more complexity compared to the electric field, highlighting intricate electromagnetic interactions within the structure. Areas with high concentrations of electric fields, magnetic fields, and surface currents provide valuable insights into the intensity patterns of the absorber. A thorough examination of these field distributions aids in refining the design of metamaterial absorbers, offering opportunities to enhance absorption efficiency and customize their response to specific electromagnetic frequencies.

**Fig 11.**
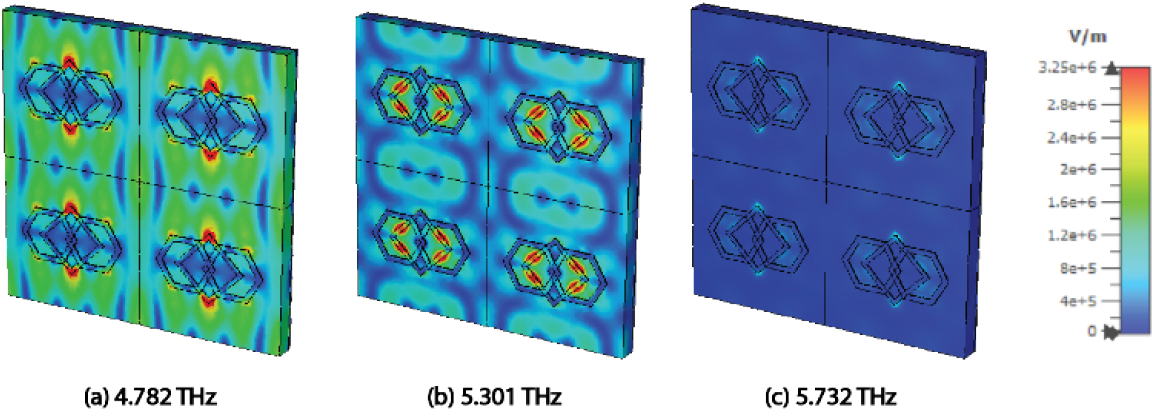
E-field distribution at (a) 4.782 THz, (b) 5.301 THz, and (c) 5.732 THz.

**Fig 12.**
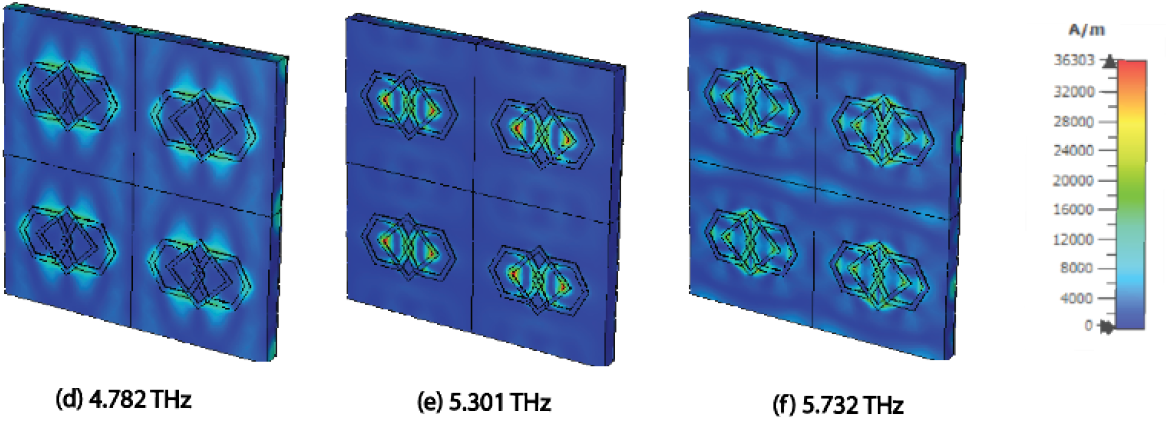
H-field distribution at (d) 4.782 THz, (e) 5.301 THz, and (f) 5.732 THz.

**Fig 13.**
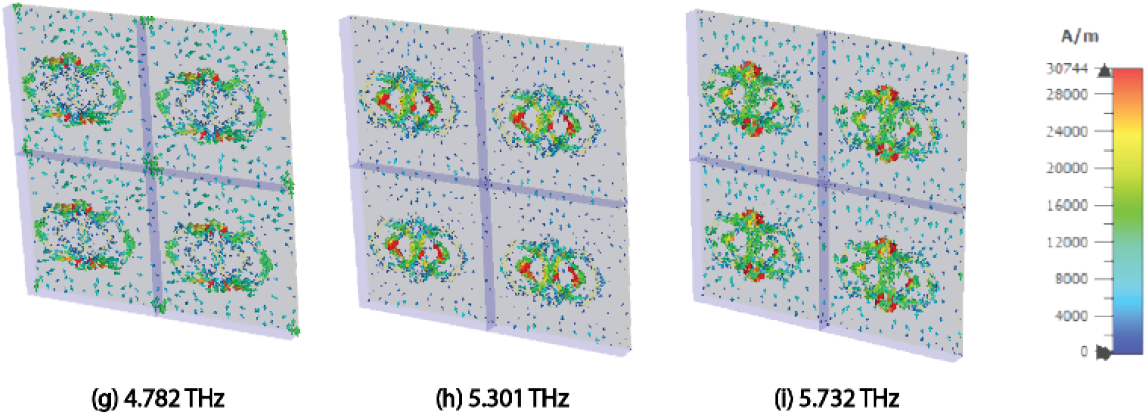
Surface current distribution at (g) 4.782 THz, (h) 5.301 THz, and (i) 5.732 THz.

**Fig 14.**
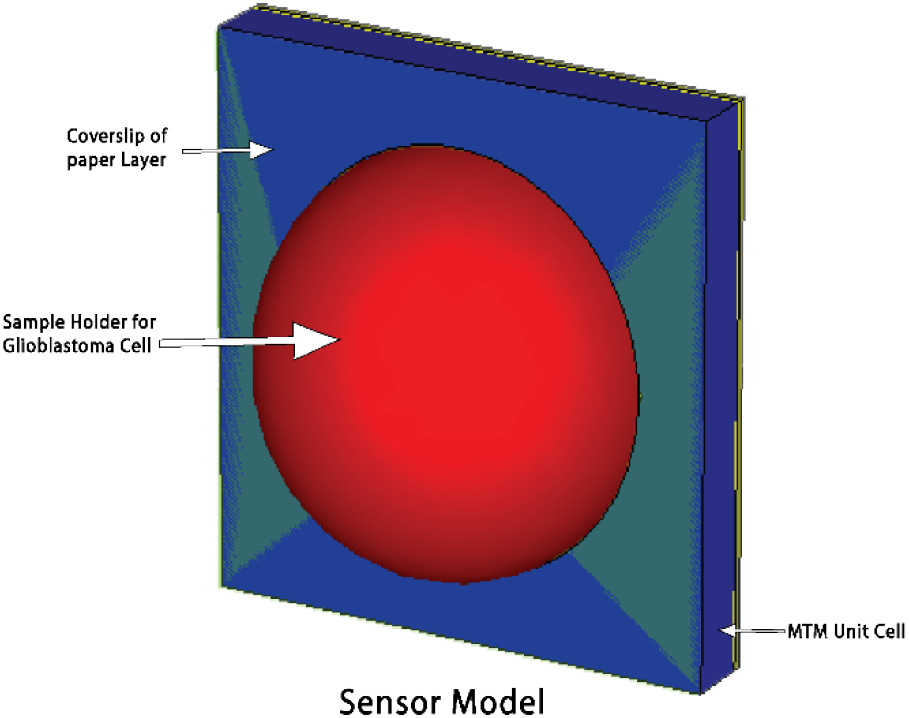
Unit cell Sensor Model.

In the fig. 11(a-c) illustrate the simulated electric field (E-field) distribution of the metamaterial absorber at three resonance frequencies—4.782 THz, 5.301 THz, and 5.732 THz—demonstrates a frequency-dependent enhancement in field concentration. At 4.782 THz, moderate E-field intensity is concentrated around the central “plus” resonator, signifying the onset of surface plasmon excitation. With an increase in frequency to 5.301 THz, the E-field extends along the hexagonal arms, indicating stronger plasmonic interactions. At the highest frequency of 5.732 THz, the field is highly localized around both the “plus” resonator and hexagonal structures, signifying the excitation of localized surface plasmon resonances (LSPR) [19]. Additionally, the substrate region between the “plus” and hexagonal elements exhibits cavity surface plasmon resonances (CSPR), illustrating the absorber’s tunability and ability to control electromagnetic fields effectively in the terahertz range.

In the fig. 11(d-f) shows the simulated magnetic field (H-field) distribution, depicted through its intensity at the same resonant frequencies, reveals frequency-dependent magnetic interactions. At 4.782 THz, moderate H-field concentration around the “plus” resonator suggests initial magnetic resonance excitation, dominated by first-order surface plasmon resonance (SPR) [20] and Fabry-Perot Resonance (FPR) [21] due to wave reflections within the unit cell. As the frequency increases to 5.301 THz, the H-field becomes more confined within the resonator structure due to Mie resonance, with second-order SPR contributing to magnetic energy distribution within the dielectric layer. At 5.732 THz, the H-field reaches its maximum localization around the “plus” resonator and hexagonal arms, reflecting strong magnetic resonance coupling and cavity magnetic resonances in the substrate. This behavior underscores the absorber’s frequency-dependent tunability, governed by the interplay of SPR, LSPR, CSPR, Mie resonance, and FPR.

The surface current distribution is depicted in Fig. 11(g-i). The graph also showcases a metasurface structure comprising multiple resonating elements. Significant coupling occurs between these elements at specific frequencies: 4.782 THz, 5.301 THz, and 5.7319 THz. This coupling is confirmed by the field distributions at these frequencies, which reveal energy concentration in specific regions. Furthermore, cross-coupling is observed between different frequency modes, notably between the 4.782 THz, 5.30 THz, and 5.7319 THz modes. The surface current analysis at the resonant frequencies further elucidates the absorber’s electromagnetic response. At 4.782 THz, dense antiparallel current flows around the resonator structure generate a strong magnetic response, evident from the concentrated H-field at the unit cell’s center. At 5.301 THz, similar circular current patterns persist, leading to enhanced magnetic field localization. At 5.732 THz, while the current circulation remains, the H-field intensity is slightly reduced, reflecting the evolution of resonant modes with frequency. These antiparallel currents confirm the presence of magnetic dipole excitation, significantly influencing the absorber’s performance by modulating field enhancement and absorption characteristics.

Glioblastoma is the most aggressive primary brain tumor whose current standard treatment associates maximal safe surgical resection, followed by radiotherapy (RT) and Temozolomide (TMZ) based chemotherapy combination. Nonetheless, despite aggressive treatment, glioblastoma is not curable [8]. All patients present a recurrence that mostly occurs close to the surgical cavity and within the volume targeted by radiation therapy. Median survival is 15 months. Such an outcome is due both to glioblastoma propensity for infiltration, degrading surgical resection efficiency, and extensive tumoral heterogeneity resulting in variable biological responses to radio-chemotherapy. Despite being a minor population of cancer cells, the cancer stem cells (CSCs) that are identified in glioblastoma (GSCs) are thought to be the major driving force behind glioblastoma biological heterogeneity and are likely to explain the high rates of glioblastoma recurrence. Indeed, GSCs are particularly aggressive and radio-resistant. Moreover, they have the ability to self-renew rapidly in vitro to form neuro spheres, and show great plasticity which could explain the high heterogeneity of glioblastoma and GBM recurrence after radiotherapy. All glioblastomas contain CSC, but unfortunately the preferential localization of these cells in the tumor volume in vivo is unknown due to the absence of studies on the sensitivity of medical images to highlight this specific cell population.

The proposed MTM sensor includes a sample container for Glaioblastoma cells. The malignant cells are extracted from the patient’s blood. Once the cells are placed in the sample container, the sensor can differentiate between normal cells and Glaioblastoma cells by analyzing their refractive index. This is done by observing a slight shift in the resonant peak of the absorption graph from its initial value. The refractive indexes (n) of normal and Glaioblastoma cells are 1.368 and 1.392, respectively. By examining the changes in frequency in each spectral range, it is possible to differentiate Glaioblastoma cells from healthy cells. By examining the changes in frequency in each spectral range, it is possible to differentiate cancerous cells from healthy cells. Glaioblastoma cells possess a greater refractive index compared to healthy cells. This leads to variations in the intensity of electric and magnetic fields. Glaioblastoma cells exhibit greater sensitivity to the electrical and magnetic forces produced by healthy cells. By evaluating the intensity of the E and H fields, it is possible to distinguish healthy cells from malignant cells. A coverslip of paper material is placed between the sample holder and the MTM absorber to obtain error-free results. Figs. a and b illustrate the simulation setup of the suggested MTM absorber-based sensor and Fig c illustrates the experimental setup for detecting Glaioblastoma cells. In this scenario, the THz source produces a 4–6 THz plane wave as it interacts with the suggested sensor. The absorption spectrum is derived from the reflected wave detected by the spectrometric apparatus. An amplifier enhances the signal and transmits it to a computer, which analyses and predicts the refractive index of the sample cells by examining the changes in the absorption spectrum. After placing the cells in the sample holder, a transverse electric wave was used to detect the differences in the absorption curve between healthy and Glaioblastoma cells.

### Diagnosis of healthy and cancerous cell

The absorption spectra presented demonstrate a distinct leftward (red) shift in the resonance frequencies of glioblastoma cells compared to normal cells, indicating a significant difference in their electromagnetic response within the 4.0–6.0 THz range. This spectral displacement suggests that the glioblastoma cells exhibit higher dielectric losses or altered permittivity, likely due to changes in cellular composition, such as increased water content and molecular density associated with malignancy. In contrast, the normal cells show resonance peaks positioned towards higher frequencies (rightward), reflecting their relatively stable and homogeneous structure. These frequency shifts, alongside variations in absorption intensity, confirm the sensitivity and specificity of the proposed terahertz metamaterial absorber in distinguishing between healthy and cancerous tissues. Such differentiation aligns with findings in recent terahertz biosensing studies, underscoring its potential for non-invasive cancer detection and biomedical diagnostics.

Terahertz (THz) imaging enables the detection of brain cancer cells by leveraging distinct absorption spectral differences between healthy and malignant tumor cells. This study explores THz imaging by analyzing the dielectric properties of healthy and glioblastoma multiforme (GBM) cells in the cerebrum. GBM, the most aggressive primary brain tumor, predominantly arises in the cerebral hemispheres, particularly within the frontal, temporal, or parietal lobes. Collected GBM cells are positioned in the sample holder of a metamaterial (MTM) sensor. Malignant cells exhibit a higher refractive index than healthy cells, resulting in measurable variations in electric (E) and magnetic (H) field intensities. Cancer cells demonstrate heightened sensitivity to electromagnetic forces generated by healthy cells, allowing differentiation between normal and malignant cells through E- and H-field analysis [58].

The proposed biosensor employs THz-based microwave imaging (MWI) to detect refractive index variations via frequency-domain data, which provides spatial information about the target. When a THz wave interacts with a material, phase shifts occur due to refractive index differences, enabling the reconstruction of internal structural details. By analyzing these phase variations, the spatial distribution of refractive index changes can be mapped, facilitating image generation for diagnostic purposes [58]. This MWI-based approach distinguishes healthy from malignant GBM cells, supporting early brain cancer detection.

The unit cell absorber demonstrates three absorption peaks at 4.782 THz, 5.30 THz, and 5.7319 THz. Figure 15 illustrates the E-field distribution for healthy and malignant GBM cells at 4.782 THz. While healthy or normal cells exhibit negligible electric fields (Fig. 20), malignant GBM cells produce intense E-field concentrations, indicated by bright red regions in the sample holder (Fig. 20). Similarly, at 5.30 THz (Fig. 16), low E-field density corresponds to healthy or normal cells (Fig. 16), whereas elevated E-field density signifies malignancy (Fig. 16). A comparable pattern is observed at 5.7319 THz (Fig. 17), where reduced E-field intensity indicates healthy or normal cells (Fig. 17), and high E-field density confirms malignant GBM presence (Fig. 17).

**Fig 15.**
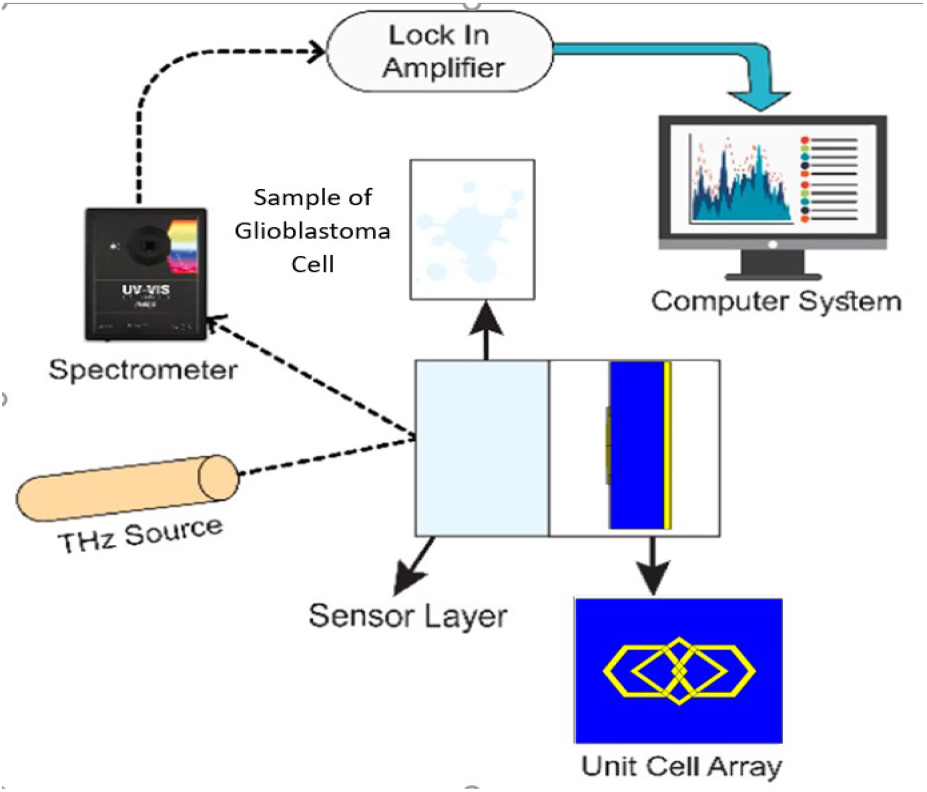
Experimental Setup.

**Fig 16.**
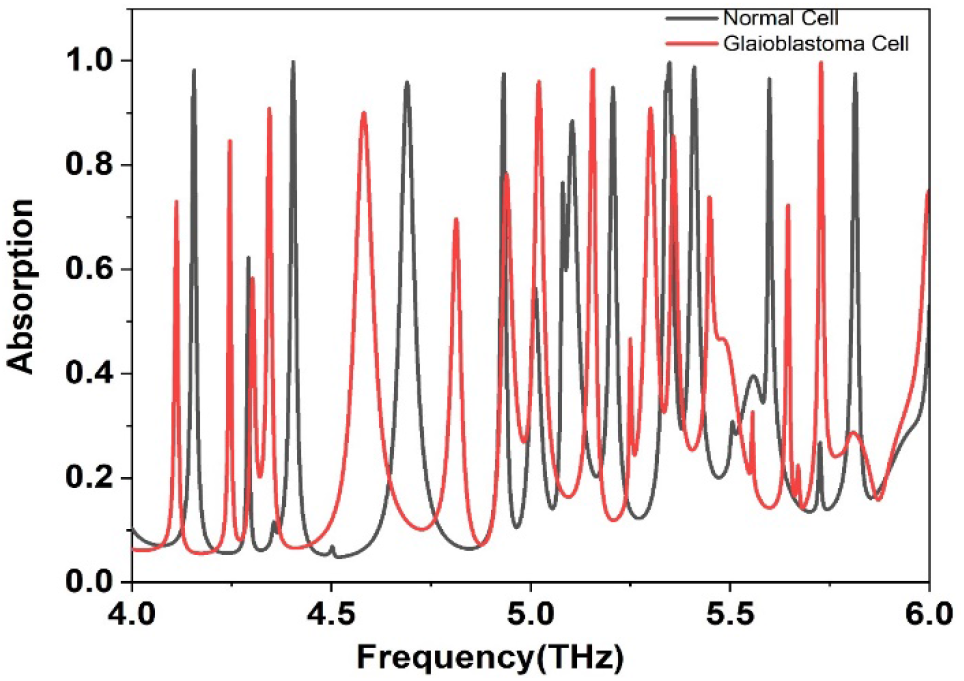
Detection of Glioblastoma cell using MTM biosensor.

**Fig 17.**
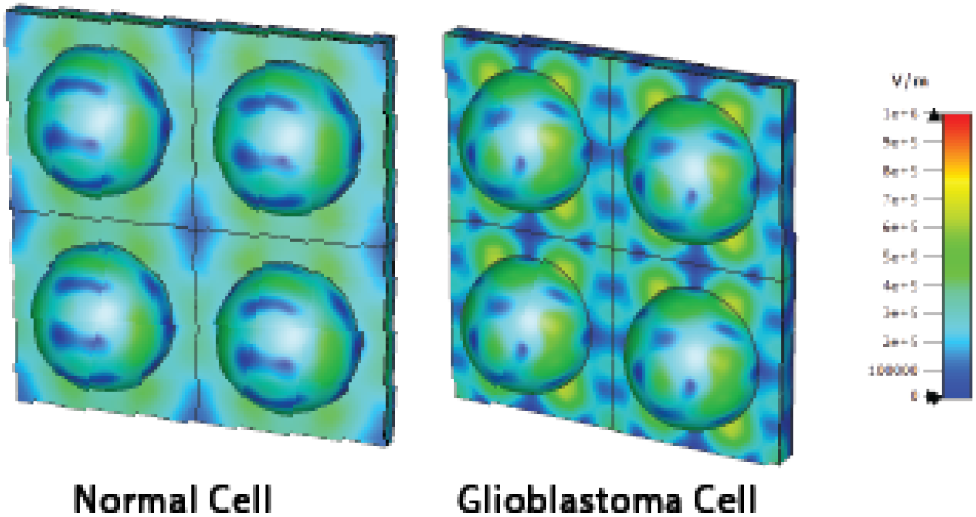
The E-field MWI result at 4.782 THz of Normal GBM cells, and cancerous GBM cells.

Further validation was conducted through magnetic field analysis. Figure 18 depicts H-field distributions at 4.782 THz, revealing low-intensity fields for normal or healthy cells (Fig. 18) and high-intensity fields for cancerous GBM cells (Fig. 18). Figure 19 shows the H-field distributions at 5.30 THz, revealing high-intensity fields for normal or healthy cells (Fig. 19) and low-intensity fields for cancerous GBM cells (Fig. 19). Additional assessments at 5.7319 THz (Fig. 20) corroborate these findings, with low H-field regions corresponding to normal or healthy cells (Fig. 20) and high H-field regions marking cancerous cells (Fig. 20).

**Fig 18.**
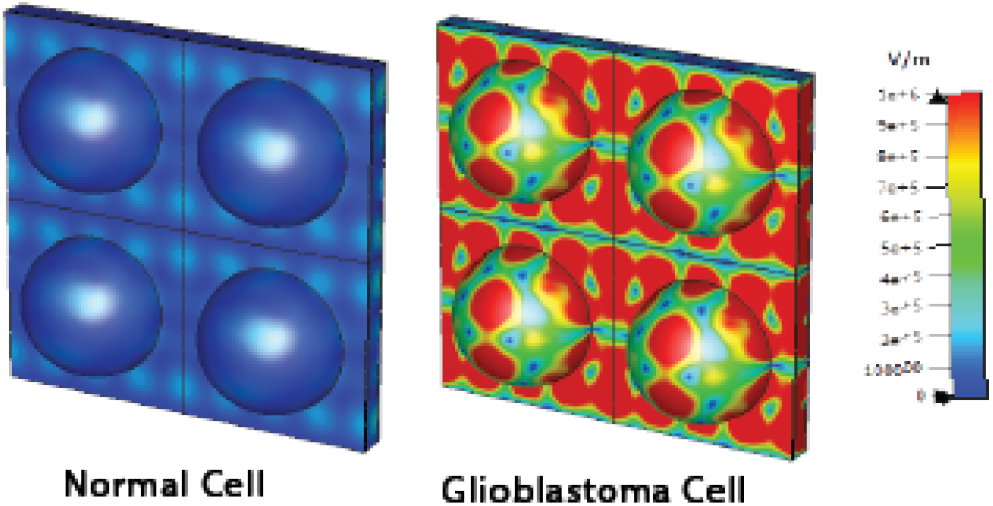
The E-field MWI result at 5.30 THz of Normal GBM cells, and cancerous GBM cells.

**Fig 19.**
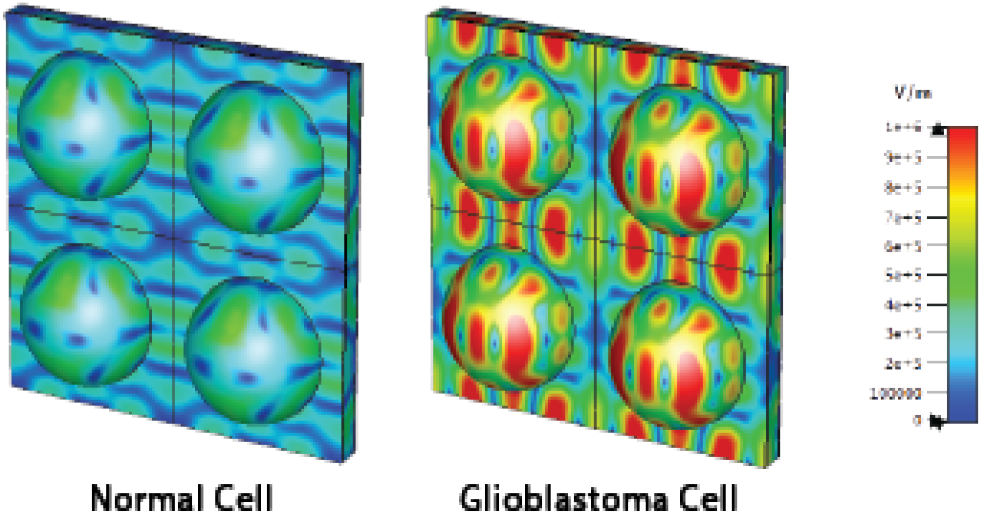
The E-field MWI result at 5.719 THz of Normal GBM cells, and cancerous GBM cells.

**Fig 20.**
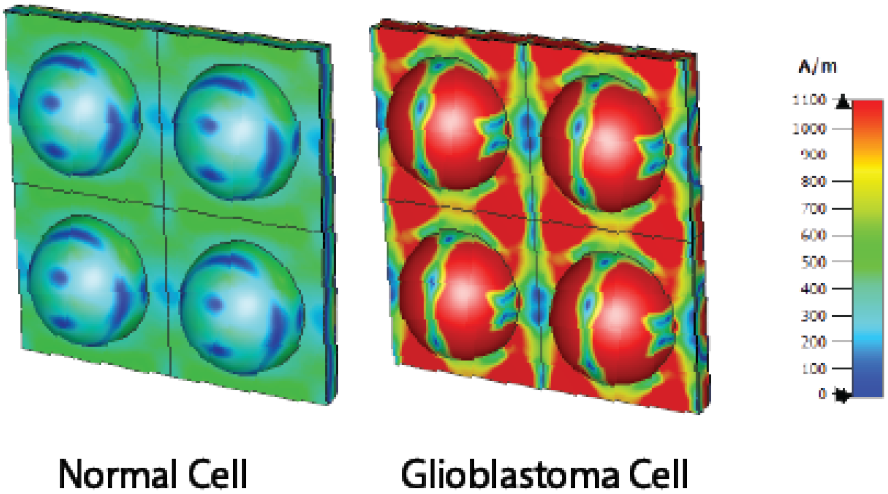
The H-field MWI result at 4.782 THz of Normal GBM cells, and cancerous GBM cells.

**Fig 21.**
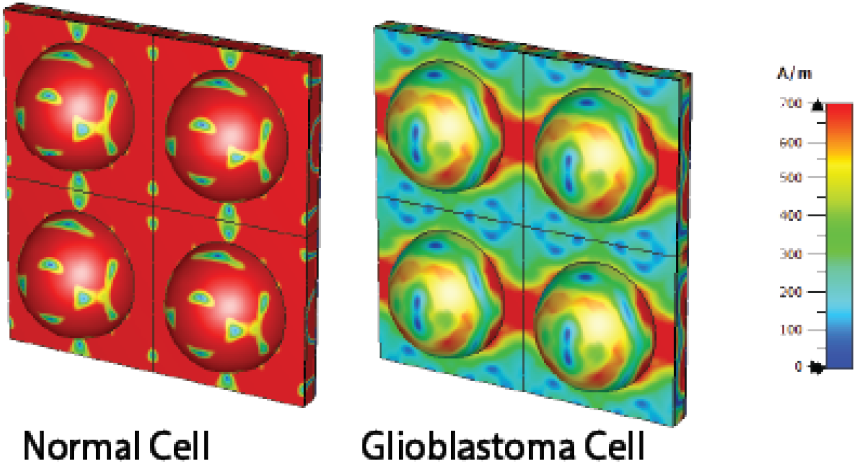
The H-field MWI result at 5.30 THz of Normal GBM cells, and cancerous GBM cells.

**Fig 22.**
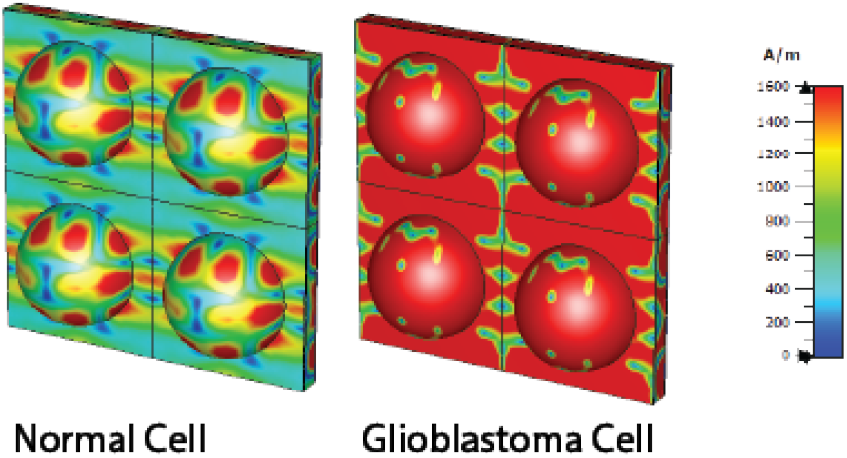
The H-field MWI result at 5.719 THz of Normal GBM cells, and cancerous GBM cells.

These results underscore the potential of magnetic field analysis as a diagnostic tool for brain cancer. The proposed metamaterial biosensor demonstrates high reliability and efficiency in detecting malignant GBM cells, offering a promising approach for early-stage cancer diagnosis and timely therapeutic intervention.

## CONCLUSION

The paper elucidates the design and analysis of a novel terahertz metamaterial absorber for detecting Glioblastoma cell. Multiple structural designs are examined, and the selected design achieves absorption rates of 99.99%, 99.98%, and 99.68% at frequencies of 4.782 THz, 5.30 THz, and 5.7319 THz, respectively. Many analyses have been examined in this paper including parametric study, quality factor, PCR, E-field, H-field, and surface current distribution. Finally, Glioblastoma cells are successfully detected using the microwave imaging method. The proposed biosensor exhibits exceptional quality factors, unique design, sensitivity, and figure of merit. With its numerous advantages, the sensor can be effectively used in biomedical applications to differentiate between Glioblastoma cells and healthy cells.

## References

[1] M. I. Hoque, N. Alam, M. K. Uddin, S. Hannan, N. M. Sahar, and A. Hoque, “Labyrinth Maze-Shaped Split Ring EM Metamaterial Absorber for C and X Band Application,” Springer Proc. Phys., vol. 303, no. July, pp. 155–164, 2024, doi: 10.1007/978-981-97-0142-1_16.

[2] D. Lee, H. Jeong, and S. Lim, “Electronically Switchable Broadband Metamaterial Absorber,” Sci. Rep., vol. 7, no. 1, pp. 1–10, 2017, doi: 10.1038/s41598-017-05330-z.

[3] H. Huang, H. Xia, W. Xie, Z. Guo, H. Li, and D. Xie, “Design of broadband graphene-metamaterial absorbers for permittivity sensing at mid-infrared regions,” Sci. Rep., vol. 8, no. 1, pp. 1–11, 2018, doi: 10.1038/s41598-018-22536-x.

[4] M. E. Mustafa, F. A. Tahir, M. Amin, and O. Siddiqui, “Comment on ‘a novel ultrathin and broadband microwave metamaterial absorber’ [J. Appl. Phys. 116, 094504 (2014)],” J. Appl. Phys., vol. 124, no. 14, Oct. 2018, doi: 10.1063/1.5046894.

[5] S. M. Sijan, S. Hannan, O. Faruk, M. Ibrahim, and M. A. Hossain, “Eye-Shaped Labyrinth Metamaterial Absorber For C, X. K. & K-Band Applications,” 5th IEEE Int. Conf. Telecommun. Photonics, ICTP 2023 - e-Proceedings, no. June, pp. 1–5, 2023, doi: 10.1109/ICTP60248.2023.10491066.

[6] M. Ahmed et al., “Analysis of a New High-Performed Terahertz Metamaterial Absorber for Refractive Index Sensor Applications,” 2024 3rd Int. Conf. Adv. Electr. Electron. Eng. ICAEEE 2024, no. June, 2024, doi: 10.1109/ICAEEE62219.2024.10561731.

[7] A. Khamsavi, Y. Abdi, Z. Fekrirad, and E. Arefian, “Graphene/Si-Based Biosensor for Glioblastoma,” vol. 22, no. 6, pp. 5548–5554, 2022.

[8] M. N. Hamza and M. T. Islam, “Designing an Extremely Tiny Dual-Band Biosensor Based on MTMs in the Terahertz Region as a Perfect Absorber for Non-Melanoma Skin Cancer Diagnostics,” IEEE Access, vol. 11, pp. 136770–136781, 2023, doi: 10.1109/ACCESS.2023.3339562.

[9] C. Tan et al., “Cancer Diagnosis Using Terahertz-Graphene-Metasurface-Based Biosensor with Dual-Resonance Response,” Nanomaterials, vol. 12, no. 21, 2022, doi: 10.3390/nano12213889.

[10] J. Yang and Y. S. Lin, “Design of tunable terahertz metamaterial sensor with single-and dual-resonance characteristic,” Nanomaterials, vol. 11, no. 9, 2021, doi: 10.3390/nano11092212.

[11] P. Jain et al., “Machine learning assisted hepta band THz metamaterial absorber for biomedical applications,” Sci. Rep., vol. 13, no. 1, pp. 1–13, 2023, doi: 10.1038/s41598-023-29024-x.

[12] Y. I. Abdulkarim et al., “Highly Sensitive Dual-Band Terahertz Metamaterial Absorber for Biomedical Applications: Simulation and Experiment,” ACS Omega, vol. 7, no. 42, pp. 38094–38104, 2022, doi: 10.1021/acsomega.2c06118.

[13] S. Asgari and T. Fabritius, “Multi-band terahertz metamaterial absorber composed of concentric square patch and ring resonator array,” Opt. Contin., vol. 3, no. 2, pp. 148–163, 2024, doi: 10.1364/OPTCON.506061.

[14] M. E. Mustafa, F. A. Tahir, M. Amin, and O. Siddiqui, “Comment on ‘a novel ultrathin and broadband microwave metamaterial absorber’ [J. Appl. Phys. 116, 094504 (2014)],” J. Appl. Phys., vol. 124, no. 14, pp. 1–5, 2018, doi: 10.1063/1.5046894.

[15] B. X. Wang, X. Zhai, G. Z. Wang, W. Q. Huang, and L. L. Wang, “Design of a four-band and polarization-insensitive terahertz metamaterial absorber,” IEEE Photonics J., vol. 7, no. 1, 2015, doi: 10.1109/JPHOT.2014.2381633.

[16] M. R. Foysal et al., “Design and Optimization of a Thin, Multi-Resonant Metamaterial Absorber for Enhanced Electromagnetic Absorption in C and X Bands,” pp. 4–9, 2000.

[17] S. J. Park, S. H. Cha, G. A. Shin, and Y. H. Ahn, “Sensing viruses using terahertz nano-gap metamaterials,” Biomed. Opt. Express, vol. 8, no. 8, pp. 3551, 2017, doi: 10.1364/boe.8.003551.

[18] T. Chen, D. Zhang, F. Huang, Z. Li, and F. Hu, “Design of a terahertz metamaterial sensor based on split ring resonator nested square ring resonator,” Mater. Res. Express, vol. 7, no. 9, pp. 95802, 2020, doi: 10.1088/2053-1591/abb496.

[19] A. Miah et al., “Double negative (DNG) square-enclosed C.S.R.R. metamaterial for liquid sensing applications with high sensitivity,” 2024 3rd Int. Conf. Adv. Electr. Electron. Eng. ICAEEE 2024, no. June, 2024, doi: 10.1109/ICAEEE62219.2024.10561846.

[20] C.-H. Fann, J. Zhang, M. ElKabbash, W. R. Donaldson, E. Michael Campbell, and C. Guo, “Broadband infrared plasmonic metamaterial absorber with multipronged absorption mechanisms,” Opt. Express, vol. 27, no. 20, pp. 27917, 2019, doi: 10.1364/oe.27.027917.

[21] S. M. A. Haque, M. Ahmed, A. Alqahtani, M. R. Maruf, M. T. Islam, and M. Samsuzzaman, “Modelling and analysis of a triple-band metamaterial absorber for early-stage cervical cancer HeLa cell detection,” Opt. Lasers Eng., vol. 181, no. June, p. 108426, 2024, doi: 10.1016/j.optlaseng.2024.108426.

[22] R. M. H. Bilal et al., “Elliptical metallic rings-shaped fractal metamaterial absorber in the visible regime,” Sci. Rep., vol. 10, no. 1, pp. 1–13, 2020, doi: 10.1038/s41598-020-71032-8.

[23] H. Hu, B. Qi, Y. Zhao, X. Zhang, Y. Wang, and X. Huang, “A graphene-based THz metasurface sensor with air-spaced structure,” Front. Phys., vol. 10, no. September, pp. 1–11, 2022, doi: 10.3389/fphy.2022.990126.

[24] R. Bhati and A. K. Malik, “Multiband terahertz metamaterial perfect absorber for microorganisms detection,” Sci. Rep., vol. 13, no. 1, pp. 1–12, 2023, doi: 10.1038/s41598-023-46787-5.

